# Chronic Pain Patients Exhibit Individually Unique Cortical Signatures of Pain

**DOI:** 10.1101/2020.09.05.284117

**Authors:** Astrid Mayr, Pauline Jahn, Anne Stankewitz, Bettina Deak, Anderson Winkler, Viktor Witkovsky, Ozan Eren, Andreas Straube, Enrico Schulz

## Abstract

We investigated how the trajectory of pain patients’ ongoing and fluctuating pain is encoded in the brain. In repeated fMRI sessions, 20 chronic back pain patients and 20 chronic migraineurs were asked to continuously rate the intensity of their endogenous pain. Linear mixed effects models were used to disentangle cortical processes related to pain intensity and to pain intensity changes. We found that the intensity of pain in chronic back pain patients is encoded in the anterior insula, the frontal operculum, and the pons; the change of pain of chronic back pain and chronic migraine patients is mainly encoded in the anterior insula. At the individual level, we identified a more complex picture where each patient exhibited their own signature of endogenous pain encoding. The diversity of the individual cortical signatures of chronic pain encoding results adds to the understanding of chronic pain as a complex and multifaceted disease.

## Introduction

The perception of pain is a subjective and multidimensional experience that has a profound impact on the physiological and psychological state of an individual (Baliki et al., 2006; Moriarty et al., 2011). Chronic pain states are characterised by a hypersensitisation of nociceptive neurons (Gold and Gebhart, 2010), a reduced endogenous inhibition of the nociceptive system, and by maladaptive cortical processes (Phillips, 2009; Price, 2000). The neural mechanisms that underlie the process of chronification are poorly understood. Several studies found the amygdala, the medial prefrontal cortex, and the nucleus accumbens to be important in the transition from subacute to chronic back pain (Hashmi et al., 2013; Makary et al., 2020). Additionally, the activity of the posterior hypothalamus is suggested to differentiate between episodic and chronic migraineurs (Schulte et al., 2017).

The cortical regions that are involved in the processing of chronic pain have primarily been investigated with experimentally-applied exogenous pain, i.e. thermal (Baliki et al., 2010; Gollub et al., 2018; Schwedt et al., 2011; Vachon-Presseau et al., 2016), electrical (Callan et al., 2014; Diers et al., 2007; Lloyd et al., 2008), mechanical (Giesecke et al., 2004; Gracely et al., 2002; Grossi et al., 2011; Kobayashi et al., 2009), or chemical stimulation (Schulte et al., 2017; Stankewitz et al., 2010). These studies found increased activity in chronic pain patients in regions already known to be involved in the encoding of acute pain, such as the primary and secondary somatosensory cortices (S1, S2), sections of the insular and cingulate cortices, the cerebellum, and the thalamus (Apkarian et al., 2005; Baliki et al., 2006; Filippi and Messina, 2020; Malinen et al., 2010; Price, 2000). However, applying experimental pain to patients and investigating pain-unspecific cortical networks may reveal cortical processes that are not necessarily at the core of the pain disease.

What matters to the individual, however, are the dynamics of the endogenous pain experience, which fluctuates over time and consists of periods of increasing, stable, and decreasing pain. To ameliorate the suffering, research should focus on cortical processes that reflect the processing of a patient’s own endogenous pain. A number of studies have utilised this experimental approach with CBP patients continuously evaluating their spontaneously fluctuating pain without external stimulation (Baliki et al., 2006; Hashmi et al., 2012a, 2012b; May et al., 2019). Findings have reported higher activity in the medial prefrontal cortex for high pain compared to low pain intensity.

Similarly, larger amplitudes of neuronal gamma oscillations have been recorded for higher pain intensities at frontocentral electrode sites in EEG (May et al., 2019). When contrasting periods of increasing pain to periods of stable and decreasing pain, Baliki and colleagues found an effect in the right anterior and posterior insula, S1 and S2, the middle cingulate cortex, and the cerebellum (Baliki et al., 2006). However, the investigation of pain intensity coding should not be restricted to periods of increasing pain (Baliki et al., 2011; May et al., 2019), but should also be assessed during periods of decreasing pain. Additionally, in some patients, higher pain intensities have been assessed only during the second half of the experiment, which renders it impossible to exclude any effect of order (May et al., 2019). As a methodological challenge in continuous and event-free *imaging* studies, it is important to preserve the naturally evolving cortical trajectory of the patients’ endogenous pain. For the present study, using functional MRI, we took a number of measures to tackle these issues, e.g. by preserving the natural autocorrelation, by applying only moderate data filtering, and by excluding recordings with pain timecourses that do not exhibit any fluctuation.

Here, we aim at reliably assessing the neuronal underpinnings of the natural fluctuations of spontaneous pain in four repeated sessions of functional imaging of two chronic pain diseases: chronic back pain and chronic migraine. Extended and continuous recordings in repeated sessions enable us to capture balanced periods of rising, stable, and falling pain in all participants. The experimental design allows us to disentangle the cortical underpinnings of pain intensity encoding from the cortical mechanisms of intensity change detection (increasing, decreasing pain), as well as from cortical mechanisms of decision making, motor processing and changes of visual input. We hypothesise that the processing of endogenous pain yields a huge variety of individual patterns of cortical activity that are qualitatively different in each patient.

## Materials and Methods

### Participants

The study included 20 patients diagnosed with chronic back pain (CBP - 16 female; aged 44±13 years) and 20 patients with chronic migraine (CM - 18 female; aged 34±13 years). All participants gave written informed consent. The study was approved by the Ethics Committee of the Medical Department of the Ludwig-Maximilians-Universität München and conducted in conformity with the Declaration of Helsinki.

CBP patients are diagnosed according to the IASP criteria (The International Association for the Study of Pain; (Merskey and Bogduk, 1994)), which includes a disease duration of more than 6 months (mean CBP: 10±7 years). All patients were seen in a specialised pain unit. CM patients are diagnosed according to the ICHD-3 (Headache Classification Committee of the International Headache Society (IHS), 2018), defined as a headache occurring on 15 or more days/month for more than 3 months, which, on at least 8 days/month, has the features of migraine headache (mean CM: 15±12 years). All CM patients were seen in a tertiary headache centre.

All patients were permitted to continue their pharmacological treatment at a stable dose (Supplementary Table 1 and Supplementary Table 2). The patients did not report any other neurological or psychiatric disorders, or had contraindications for an MRI examination. Patients who had any additional pain were excluded. For all patients, pain was fluctuating and not constant at the same intensity level. Patients with no pain or a migraine attack on the day of the measurement were asked to return on a different day. Patients were characterised using the German Pain Questionnaire (Deutscher Schmerzfragebogen; (Casser et al., 2012)) and the German-version of the Pain Catastrophizing Scale (PCS; (Sullivan et al., 1995); Table 1). The pain intensity describes the average pain in the last 4 weeks from zero to 10 with zero representing no pain and 10 indicating maximum imaginable pain (please note that this scale differs from the one used in the fMRI experiment). The Depression, Anxiety and Stress Scale (DASS) was used to rate depressive, anxiety, and stress symptoms over the past week on three 7-item subscales using 4-point scales (Lovibond and Lovibond, 1995).

**Table 1.** Active brain areas that encode pain intensity across all CBP patients.

| Anatomical structure | Cluster size | t-value |  | Coordinates |  |  |
| --- | --- | --- | --- | --- | --- | --- |
|  |  | pos | neg | x | y | z |
| Posterior Cingulate Cortex (PCC), Precuneus | 1895 |  | -5.40 | 0 | -55 | 26 |
| Anterior Insular Cortex (AIC) | 1236 | 6.50 |  | 38 | 15 | 2 |
| Perigenual Anterior Cingulate Cortex (pACC) | 507 |  | -4.99 | -1 | 44 | -3 |
| Frontal Opercular Cortex, Anterior Insular Cortex (AIC) | 395 | 4.92 |  | -43 | 16 | 0 |
| Brain-Stem (Pons) | 332 | 4.83 |  | 4 | -21 | -37 |
| Lingual Gyrus | 275 |  | -4.52 | -11 | -80 | -5 |
| Occipital Pole | 156 |  | -4.32 | 7 | -89 | 14 |
| Hippocampus, Parahippocampal Gyrus | 138 |  | -4.71 | -27 | -38 | -12 |
| Hippocampus, Parahippocampal Gyrus | 124 |  | -4.63 | 27 | -27 | -16 |
| Middle Temporal Gyrus | 60 |  | -4.42 | -61 | -13 | -11 |
| Angular Gyrus, Lateral Occipital Gyrus | 43 |  | -3.82 | -42 | -60 | 31 |
| Occipital Fusiform Gyrus, Lingual Gyrus | 37 |  | -3.90 | 23 | -71 | -6 |
| Lateral Occipital Cortex | 34 |  | -3.69 | 56 | -65 | 24 |
| Frontal Pole | 32 | 4.17 |  | -39 | 42 | 1 |
| Frontal Pole | 30 | 3.90 |  | 38 | 42 | 11 |
| Cerebellum (Left VIIb) | 25 | 3.81 |  | -34 | -53 | -48 |
| Angular Gyrus, Lateral Occipital Gyrus | 25 |  | -3.93 | 48 | -57 | 32 |
| Frontal Pole | 24 |  | -3.66 | -41 | 59 | 2 |
| Frontal Medial Cortex | 23 |  | -3.85 | 1 | 41 | -17 |
| Cerebellum (Left IX) | 22 |  | -3.76 | -9 | -49 | -41 |
| Brainstem (Pons) | 21 | 3.74 |  | -12 | -25 | -42 |

**Table 2.** Active brain areas that encode the change of pain intensity across all CBP patients.

| Anatomical structure | Cluster size | t-value |  | Coordinates |  |  |
| --- | --- | --- | --- | --- | --- | --- |
|  |  | pos | neg | x | y | z |
| Anterior Insular Cortex | 487 | 6.19 |  | -34 | 24 | 0 |
| Anterior Insular Cortex | 182 | 5.03 |  | 33 | 29 | 2 |
| Occipital Pole | 123 |  | -5.01 | -20 | -98 | -1 |
| Frontal Pole | 109 | 4.72 |  | -39 | 48 | 16 |
| Inferior Frontal Gyrus | 102 | 4.82 |  | -53 | 10 | 9 |
| Lateral Occipital Cortex | 94 |  | -5.54 | 50 | -60 | 32 |
| Frontal Pole | 76 |  | -5.28 | 23 | 40 | 44 |
| Cerebellum (Left IX) | 50 |  | -5.34 | -9 | -51 | -49 |
| Superior Frontal Gyrus | 48 |  | -4.50 | 25 | 25 | 58 |
| Inferior Frontal Gyrus, Frontal Pole | 46 | 4.62 |  | -48 | 34 | 14 |
| Posterior Cingulate Cortex (PCC) | 24 | 4.40 |  | -2 | -26 | 26 |
| Perigenual Anterior Cingulate Cortex (pACC) | 24 |  | -4.42 | 3 | 38 | -8 |

Patients were compensated with 60€ for each session. In total nine screened patients were excluded: two patients developed additional pain during the study, the pain ratings of five patients were constantly increasing or decreasing throughout the pain rating experiment, and two patients were unable to comply with study requests. 36 patients were recorded four times across 6 weeks with a gap of at least 2 days (CBP = 9±12 days, CM = 12±19 days) between sessions. Four patients (2 CBP and 2 CM) were recorded three times.

### Experimental procedure

During the recording of fMRI, patients rated the intensity of their ongoing pain for 25 minutes using an MRI-compatible potentiometer slider (Schulz et al., 2019). The scale ranged from zero to 100 in steps of five with zero representing no pain and 100 representing the highest experienced pain. On a screen a moving red cursor on a dark grey bar (visual analogue scale, VAS) and a number above (numeric analogue scale, NAS) were shown during the entire functional MRI session. The screen was visible through a mirror mounted on top of the MRI head coil. Patients were asked to look only at the screen, focus on their pain with an emphasis on rising and falling pain. The intensity and the changes of perceived pain had to be indicated as quickly and accurately as possible. To minimise head movement, foams were placed around the head and patients were told to lie as still as possible.

To control for visual-motor performance and decision-making activity during the pain rating session, a visual control experiment was carried out during the last visit. Patients were asked to continuously rate the changing background brightness of the screen as accurately and quickly as possible. The same feedback in the form of a red bar (VAS) and a number (NAS) was given on the same screen as for the pain-rating experiment. Unknown to the patient, the control condition was a composition of parts of the previous pain rating sessions to match with range and frequency of the pain rating. Pain ratings between 0 and 100 in steps of 5 were converted to 21 shades of grey, 0 indicating black ([RGB: 0 0 0]) and 100 indicating white ([RGB: 255 255 255]).

### Data Acquisition

Data were recorded on a clinical 3 tesla MRI scanner (Siemens Magnetom Skyra, Germany) using a 64-channel head coil. A T2*-weighted BOLD (blood oxygenation level dependent) gradient echo sequence with echo-planar image acquisition and a multiband factor of 2 was used with the following parameters: number of slices = 46; repetition time/echo time = 1550/30 ms; flip angle = 71°; slice thickness = 3 mm; voxel size = 3×3×3 mm^3^; field of view = 210 mm^2^. 1000 volumes were recorded in 1550 seconds. Field maps were acquired in each session to control for B0-effects. For each patient, T1-and T2-weighted anatomical MRI images were acquired using the following parameters for T1: repetition time/echo time = 2060/2.17 ms; flip angle = 12°; number of slices = 256; slice thickness = 0.75 mm; field of view = 240×240 mm^2^, and for T2: repetition time/echo time = 3200/560 ms; flip angle = 120°; number of slices = 256; slice thickness = 0.75 mm; field of view = 240×240 mm^2^.

### Data processing - behavioural data

The rating data were continuously recorded with a variable sampling rate but downsampled offline at 10 Hz. To remove the same filtering effects from the behavioural data as from the imaging data, we applied a 400 s high-pass filter (see below). For the statistical analysis, the resulting filtered timecourse was transferred to Matlab (Mathworks, USA; version R2018a) and downsampled to the sampling frequency of the imaging data (1/1.55 Hz). We shifted the rating vector between −5 s and 15 s in steps of 0.5 s (41 steps). These systematic shifts would account for (a) the unknown delay of the BOLD response and (b) for the unknown timing of cortical processing in reference to the rating: some ongoing cortical processes may influence later changes in pain ratings, other processes are directly related to the rating behaviour, or are influenced by the rating process and are occurring afterwards. We are aware that the variable timing of the BOLD response and the variable timing of the cortical processes are intermingled and would not interpret the timing aspects any further.

To disentangle the distinct aspects of pain intensity (AMP - amplitude) from cortical processes related to the sensing of rising and falling pain, we generated a further vector by computing the ongoing rate of change (SLP - slope, encoded as 1, −1, and 0) in the pain ratings. The rate of change is represented as the slope of the regression of the least squares line across a 3 s time window of the 10 Hz pain rating data. We applied the same shifting (41 steps) as for the amplitude timecourse. A vector of the absolute slope of pain ratings (aSLP - absolute slope, encoded as 0 and 1), represents periods of motor activity (slider movement), changes of visual input (each slider movement changes the screen), and decision-making (each slider movement prerequisites a decision to move).

To avoid any effects of order (in case of continuously rising pain) in the data, the patients’ rating timecourses were required to fluctuate at a relatively constant level. To ensure the behavioural task performance of the patients fulfilled this criterion, the ratings of each patient’s pain was measured with a constructed parameter PR defined as follows, with “ pain_ud_/ time” being the slope of the regression line of the unfiltered data (*ud*) and σ_fd_ being the sample standard deviation of the filtered data (*fd*):

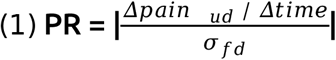

The parameter PR is constructed in a way that a minimisation of the quotient is desirable. The numerator describes how much the prerequisite is violated by fitting a least squares line across the rating time course. This violation can be compensated if the variance of pain ratings is at least four times higher than the slope of the regression of the least squares line across the entire time course of the pain ratings. The standard deviation of the filtered data expressed in the denominator gives a measure of the desired fluctuation of the pain ratings but is stripped from a potential trend throughout the experiment. A minimisation of the quotient is given either by minimising the numerator corresponding to a small increase of the overall pain ratings over the whole experiment (small slope), or by maximising the denominator corresponding to a large variability in pain ratings across the rating task. Minor overall rising in pain intensity over the whole time of the experiment could be compensated by a greater variance of ratings; small fluctuation of pain intensity would only be accepted in cases of minor pain rating trends across the entire experiment. Recordings showing PR values of ≥0.25 were rejected from the analysis or repeated if possible. We excluded five participants and repeated three sessions that exhibited a continuous rise of pain intensity throughout the experiment. For examples see Fig. 1.

**Figure 1.**
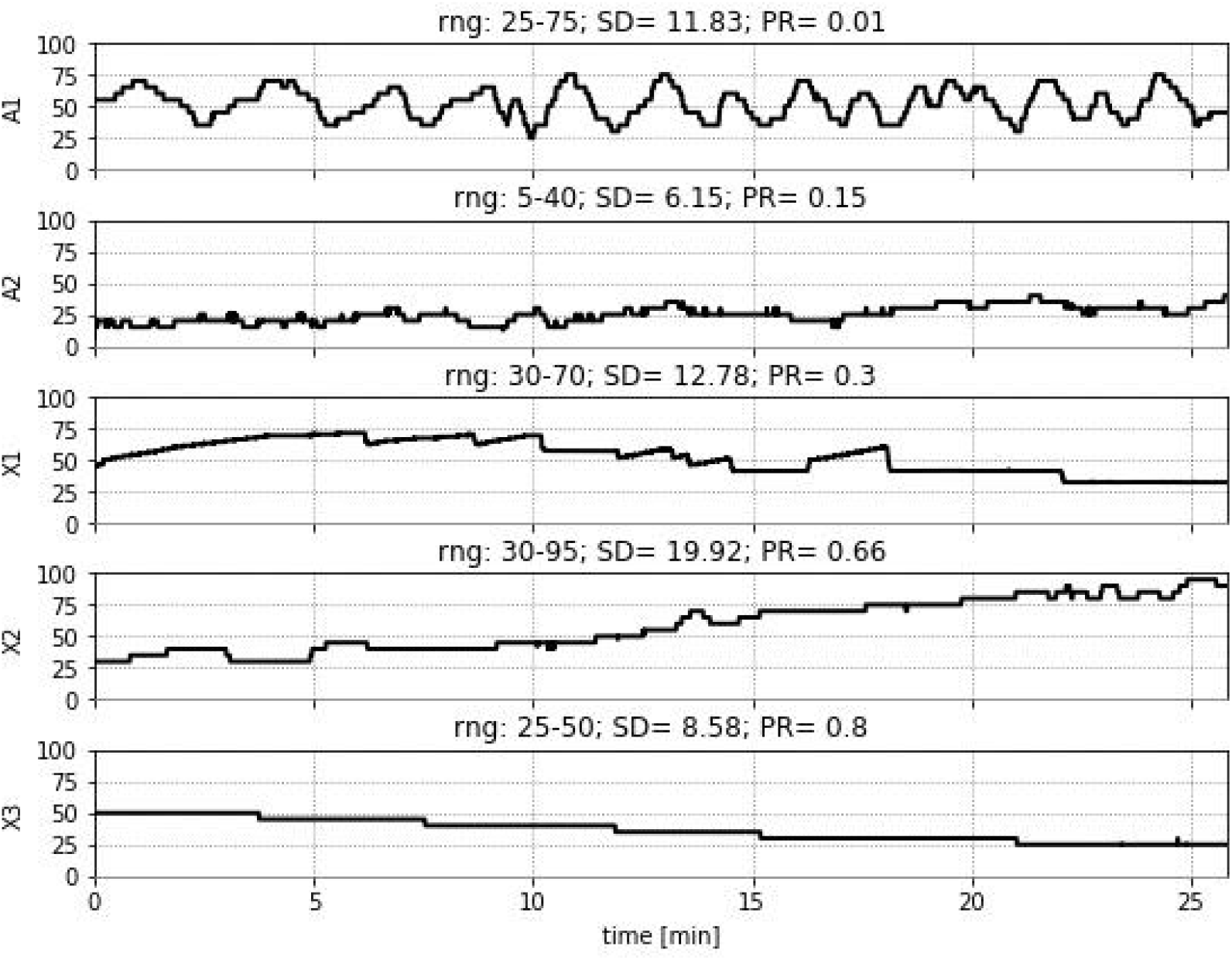
Accepted and rejected pain ratings based on the parameter PR. Rating A1 represents an excellent rating with a very low PR value of 0.01, indicating high variance and no overall drift throughout the experiment. Rating A2 represents a pain rating with a moderate PR value of 0.15, whereas X1 represents an excluded rating with a PR slightly higher than the threshold PR ≥ X2 was excluded due to a steady increase in the pain ratings over the course of the experiment. X3 was excluded due to very low variability in ratings after high-pass filtering. Ratings A1 and A2 were accepted, whereas recordings X1, X2 and X3 were excluded from the analysis.

### Data processing - imaging data

Functional MRI data were preprocessed using FSL (Version 5.0.10, (Jenkinson et al., 2012)), which included removal of non-brain data (using brain extraction), slice time correction, head motion correction, B0 unwarping, spatial smoothing using a Gaussian kernel of FWHM (full width at half maximum) 6 mm, a nonlinear high-pass temporal filtering with a cutoff of 400 s, and spatial registration to the Montreal Neurological Institute (MNI) template. The data were further semi-automatically cleaned of artefacts with MELODIC (Salimi-Khorshidi et al., 2014). Artefact-related components were evaluated according to their spatial or temporal characteristics and were removed from the data following the recommendations in (Griffanti et al., 2014; Kelly et al., 2010). The average number of artefact components for CM was 40±6 and for CBP 49±8. We deliberately did not include any correction for autocorrelation, neither for the processing of the imaging data nor for the processing of the pain rating timecourse as this step has the potential to destroy the natural evolution of the processes we aim to investigate (see PALM analysis below).

### Statistical analysis - imaging data

Using Linear Mixed Effects models (LME; MixedModels.jl package in Julia; Bezanson et al., 2015), we aimed to determine the relationship between fluctuating pain intensity and the fluctuating cortical activity separately for each voxel. The fluctuating BOLD activity of a particular brain voxel is modelled through the timecourse of the three variables (AMP, SLP, aSLP) derived from the pain ratings (Fig. 2).

**Figure 2.**
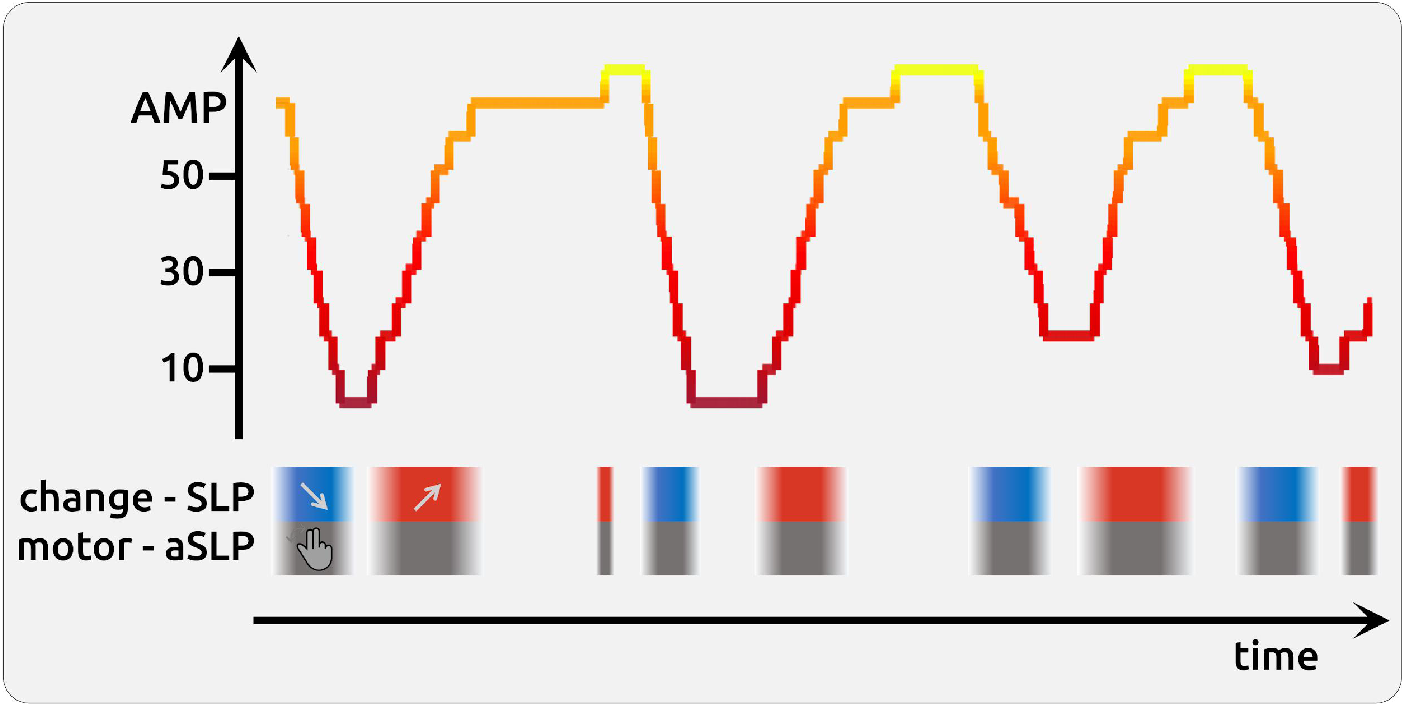
Schematic illustration of a 5 min fluctuating timecourse of pain rating. The statistical model also included the information of the direction change of pain intensity (encoded as −1 as shown in blue boxes for negative slope; encoded as 0 for stable phases of pain, and encoded as 1 as shown in red boxes for increasing pain - SLP). The task also required moving the potentiometer slider (encoded as 1 as shown in grey for motor phases, and encoded as 0 for stable phases that did not require a motor process – aSLP). Brain estimates of amplitude (AMP) should be independent irrespective of whether a data point originates from rising, stable or falling time points of the rating timecourse. In a similar vein, each slider movement involves a prior decision making and motor activity. These processes (SLP, aSLP) occur concomitant to the encoding of pain intensity (AMP) but are functionally, temporally and statistically independent.

The statistical model is expressed in Wilkinson notation; the included fixed effects (fmri ~ AMP + SLP + aSLP) describe the magnitudes of the population common intercept and the population common slopes for the relationship between cortical data and pain perception. The added random effects (e.g. AMP - 1 | session) model the specific intercept differences for each recording session (e.g. session specific differences in pain levels or echo-planar image signal intensities).

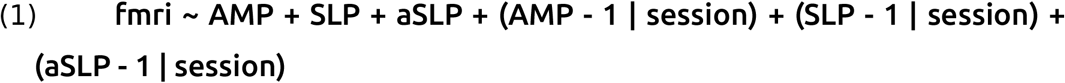

The common intercept represents the group-wise fixed effects, which are the parameters of main interest in the LME. The model also includes the random effects that represent possible differences of the effects in each session. Hence, the model has four fixed effects parameters (common intercept, AMP, SLP, aSLP) and 234 (3*78) random effect parameters with related variance components of AMP random slopes, SLP random slope, aSLP random slope, and additive unexplained error. Each model was computed 41 times along the time shift of the rating vector (−5 to 15 s in steps of 0.5 s, see above) and the highest t-values of the fixed effect parameters AMP, SLP, and aSLP were extracted.

### Correcting statistical testing - surrogate data

All voxel-wise statistical tests had to be corrected for multiple comparisons (voxels, time shifts) and autocorrelation in the behavioural data: we created 5000 surrogate timecourses using the IAAFT algorithm (Iterative Amplitude Adjusted Fourier Transform) from the original rating data, which were uncorrelated to the original rating data but had the same autocorrelation structure as the original data (Schreiber and Schmitz, 1996). Using surrogate data, the entire LME analysis was repeated 5000 times, resulting in 5000 whole-brain statistical maps for AMP, SLP and aSLP, respectively. From each map the highest absolute t-values of each repetition across the whole volume was extracted. This procedure resulted in a right-skewed distribution of 5000 values for each condition. Based on the distributions of 5000 values (for AMP, SLP, aSLP), the statistical thresholds were determined using the “palm_datapval.m” function publicly available in PALM (Winkler et al., 2016, 2014).

### Comparisons of topographies

We tested whether the topography of the amplitude encoding (AMP) resembles the topography of the neurological signature of applied *physical* pain (Neurologic Pain Signature (NPS); (Wager et al., 2013)). We thresholded the NPS weights map at 0.005 and the AMP map at t > 2 and “normalised” both maps with the following procedure: the pre-selected absolute values (5549 voxels for CM; 3324 voxels for CBP) were ranked and equidistant numbers between 1 and 1000 were given to each included voxel. Voxels with negative t-values were given back their negative sign. Spatial correlations using Kendall’s τ (tau) coefficients were computed for each patient’s activity map with the group activity map.

We also investigated the individual maps of endogenous pain encoding separately for each participant across all recordings. To assess whether the pattern of activity resembles the map of the group statistics, we correlated the activity of the group maps with the activity of the single-patient maps. The data were restricted to voxels with an absolute value of t > 2 in the group statistics. We “normalised” group map and single-patient maps with the following procedure: the pre-selected absolute t-values (121671 voxels for CM; 67719 voxels for CBP) were ranked and equidistant numbers between 1 and 1000 were given to each included voxel. Voxels with negative t-values were given back their negative sign. Separately for each patient, the analysis was further restricted to voxels for which the LME had converged. Spatial correlations using Kendall’s τ (tau) coefficients were computed for each patient’s activity map with the group activity map.

### Data availability

The data that support the findings of this study are available from the corresponding author, upon reasonable request.

## Results

### Questionnaires

The mean pain intensity specified in the questionnaires was 5±2 for CBP and 5±1 for CM. The mean duration of the chronic pain was 10±7 years for CBP and 15±12 years for CM. The scores for the PCS were 17±10 for CBP and 21±10 for CM. For CBP the depression scale was 4±3, the anxiety scale 3±2 and the stress scale 7±4. For CM the depression scale was 3±3, the anxiety scale 3±4 and the stress scale 6±4 (all results given as mean ± standard deviation). See Supplementary Table 1 and Supplementary Table 2 for detailed patient characteristics and questionnaire data.

### Behavioural data

The average pain ratings were variable between recording sessions (see Supplementary Fig. 1 and Supplementary Fig. 2 for the detailed rating timecourses of each session). For CBP and CM, we found an average rating of 39 (±14) and 40 (±15), respectively. The pain ratings within each session were fluctuating substantially, as reflected by a high variance over the 25 min of recording: σ^2^ = 109.3 (±126.6) for CBP and σ^2^ = 93.3 (±62.8) for CM. Due to the prerequisites of a low PR score, the average rating did not exhibit any systematic change over the timecourse of one experimental session. In general, we found a minimal positive slope of 0.13 (±0.41; mean change of rating unit per minute) for CBP and of −0.05 (±0.37) for CM (all mean ± standard deviation).

### Imaging results

Voxel-wise LME were computed to identify the brain regions in which the cortical activity changed during the experiment with respect to the amplitude of the pain rating (*amplitude* - AMP), as well as to rising or falling pain (*slope* - SLP). Each change of pain rating, irrespective of direction, is accompanied by motor activity, decision-making processes, and the perception of visual change on the monitor (*absolute slope* - aSLP). Respective timecourses of AMP, SLP and aSLP were related to the timecourses of brain activity in order to obtain a statistical estimate of the cortical underpinnings of these independent processes. T-values of the fixed-effects parameters quantify the relationships. The model was separately calculated for CM and CBP.

#### Encoding of pain intensity across all CBP patients (AMP)

We found the subjective intensity of endogenous pain is encoded in the anterior insular cortex (AIC), the frontal operculum, and the pons. We also found regions that exhibit decreased activity with higher pain intensities: the posterior cingulate cortex (PCC), the precuneus, the perigenual anterior cingulate cortex (pACC), and the hippocampus (Fig. 3A; Table 3). We found a spatial correlation between AMP and the NPS map of 0.41 (p > 0.001).

**Table 3.** Active brain areas that encode pain intensity across all CM patients.

| Anatomical structure | Cluster size | t-value |  | Coordinates |  |  |
| --- | --- | --- | --- | --- | --- | --- |
|  |  | pos | neg | x | y | z |
| Precuneus, Posterior Cingulate Cortex (PCC) | 126 |  | -4.02 | -1 | -47 | 8 |
| Frontal Pole | 103 |  | -4.62 | 23 | 60 | 19 |
| Lateral Occipital Cortex | 95 |  | -4.42 | 14 | -84 | 44 |
| Temporal Fusiform Cortex | 55 | 3.97 |  | 42 | -45 | -21 |
| Hippocampus | 49 |  | -3.72 | -32 | -32 | -4 |
| Perigenual Anterior Cingulate Cortex (pACC) | 46 |  | -4.04 | -0 | 30 | 1 |
| Subcallosal Cortex | 31 |  | -4.00 | -3 | 20 | -12 |
| Perigenual Anterior Cingulate Cortex (pACC) | 30 |  | -4.00 | 4 | 46 | 3 |
| Thalamus | 30 |  | -4.18 | 8 | -30 | 8 |

**Figure 3.**
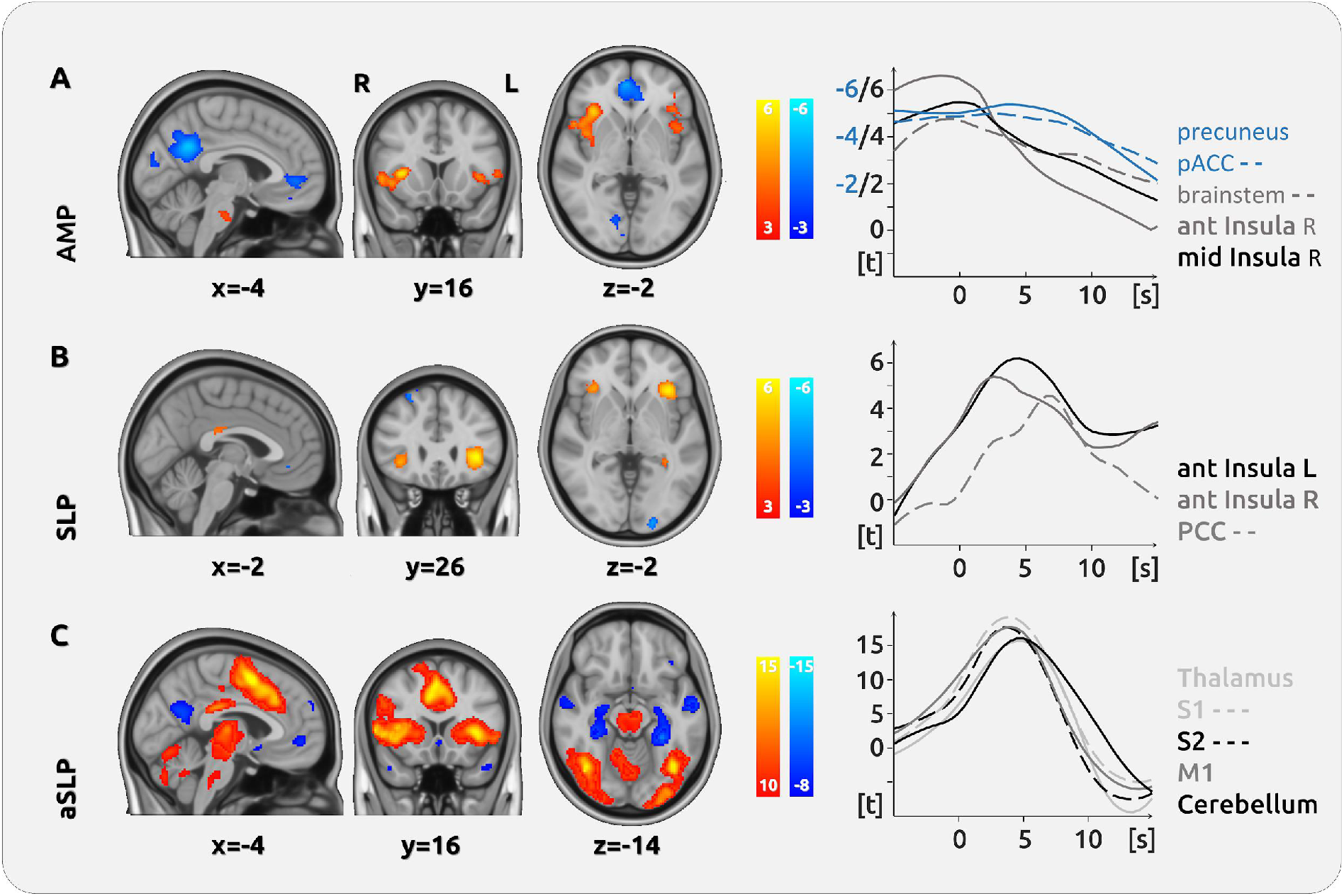
Cortical processing of chronic pain in CBP. (A) The upper row shows the cortical encoding of the endogenous pain intensity (*amplitude* - AMP): the activity in the bilateral anterior insular cortex, the pons, and the frontal cortex were positively related to pain intensity. We found negative relationships between brain activity and pain intensity in the precuneus and the perigenual ACC. (B) The processing of changes of pain intensity (*slope* - SLP) was mainly localised in the bilateral anterior insular cortex. (C) The movement of the slider entails the activation of a large network of brain regions. The movement process, which prerequisites motor activity and decision making (*absolute slope* - aSLP), shows a vast network of activity in the thalamus, the cingulate cortex, the entire insula and the cerebellum. The graphs on the right show the temporal dynamics of the haemodynamic delay for several regions in relation to a current pain rating at time point (= 0 s).

#### Encoding of the change of pain intensity across all CBP patients (SLP)

The cortical processes that reflect the change of pain intensity are located in the AIC, in various regions in the frontal lobe (frontal orbital cortex, frontal pole, inferior frontal gyrus), and in the PCC. Brain regions that exhibit decreased activity during increasing pain comprise regions in the occipital lobe (occipital pole, lateral occipital cortex), the cerebellum, regions in the frontal lobe (frontal pole, superior frontal gyrus), and the pACC (Fig. 3B; Table 4).

**Table 4.**
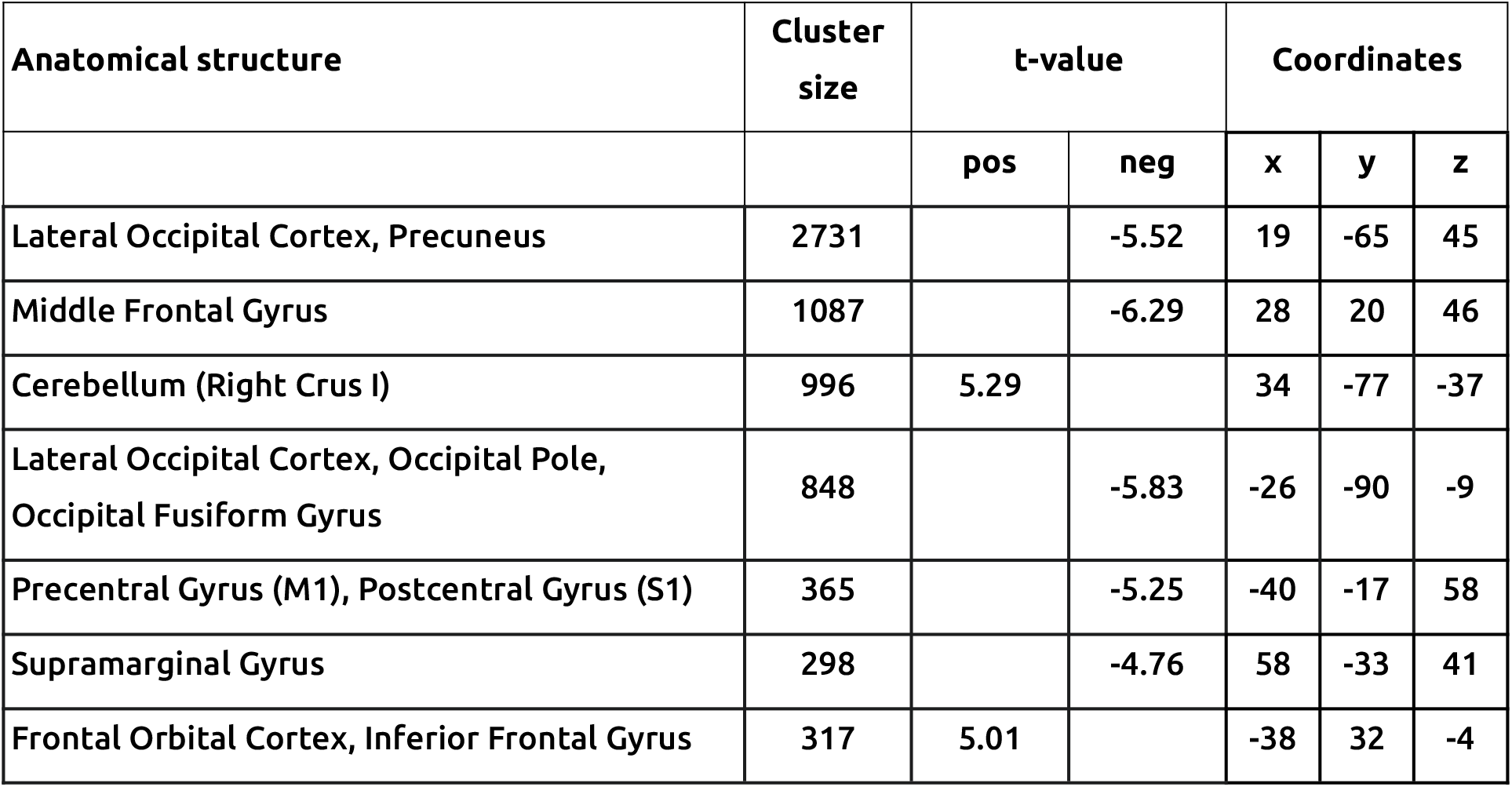
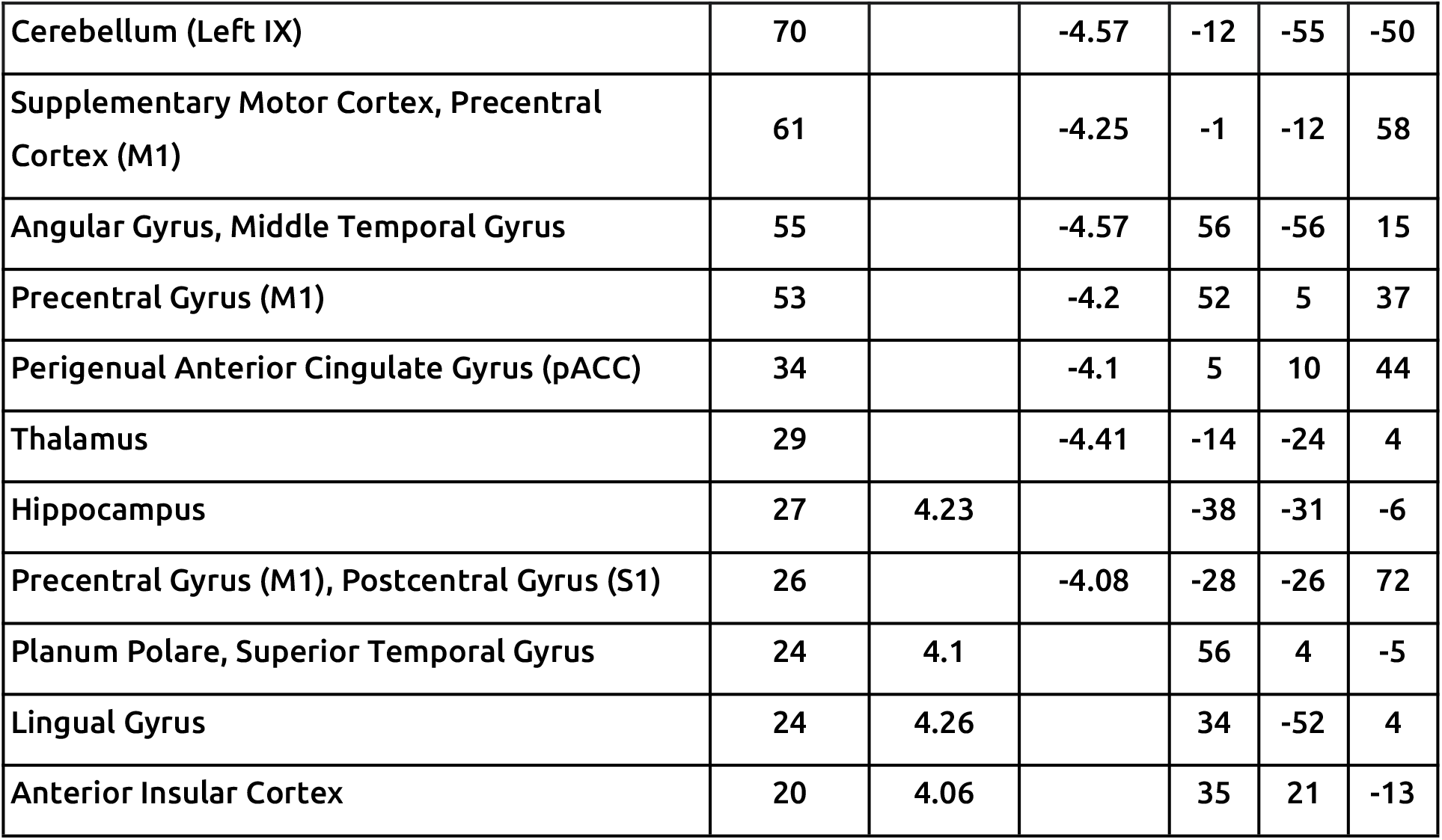
Active brain areas that encode the change of pain intensity across all CM patients.

#### Processing of motor activity and decision making across all CBP patients (aSLP)

For the processing of motor activity, decision-making, and changes of visual input, we found increased activity in the parietal lobe (superior parietal lobe, supramarginal gyrus), but also in the pre- and postcentral gyrus (M1/S1), the cerebellum, the thalamus, and the pACC. Decreased activity was located mainly in the hippocampus, the superior and middle temporal gyrus, the lateral occipital gyrus, and the precuneus. To separate the largely overlapping clusters and to disentangle the contribution of the brain regions, we increased the cluster threshold beyond the threshold of the randomisation statistics. This affects the number of activated voxels; the figure has been generated based on the original PALM threshold (Fig. 3C; Supplementary Table 3).

#### Encoding of pain intensity across all CM patients (AMP)

For CM, only the temporal fusiform cortex is positively related to the intensity of endogenous pain. We found areas that are negatively related to pain intensity in the frontal pole, the pACC, and the subcallosal cortex, but also in the lateral occipital cortex, the thalamus and the hippocampus (Fig. 4A, Table 5). We found a spatial correlation between AMP and the NPS map of −0.13 (p > 0.001).

**Figure 4.**
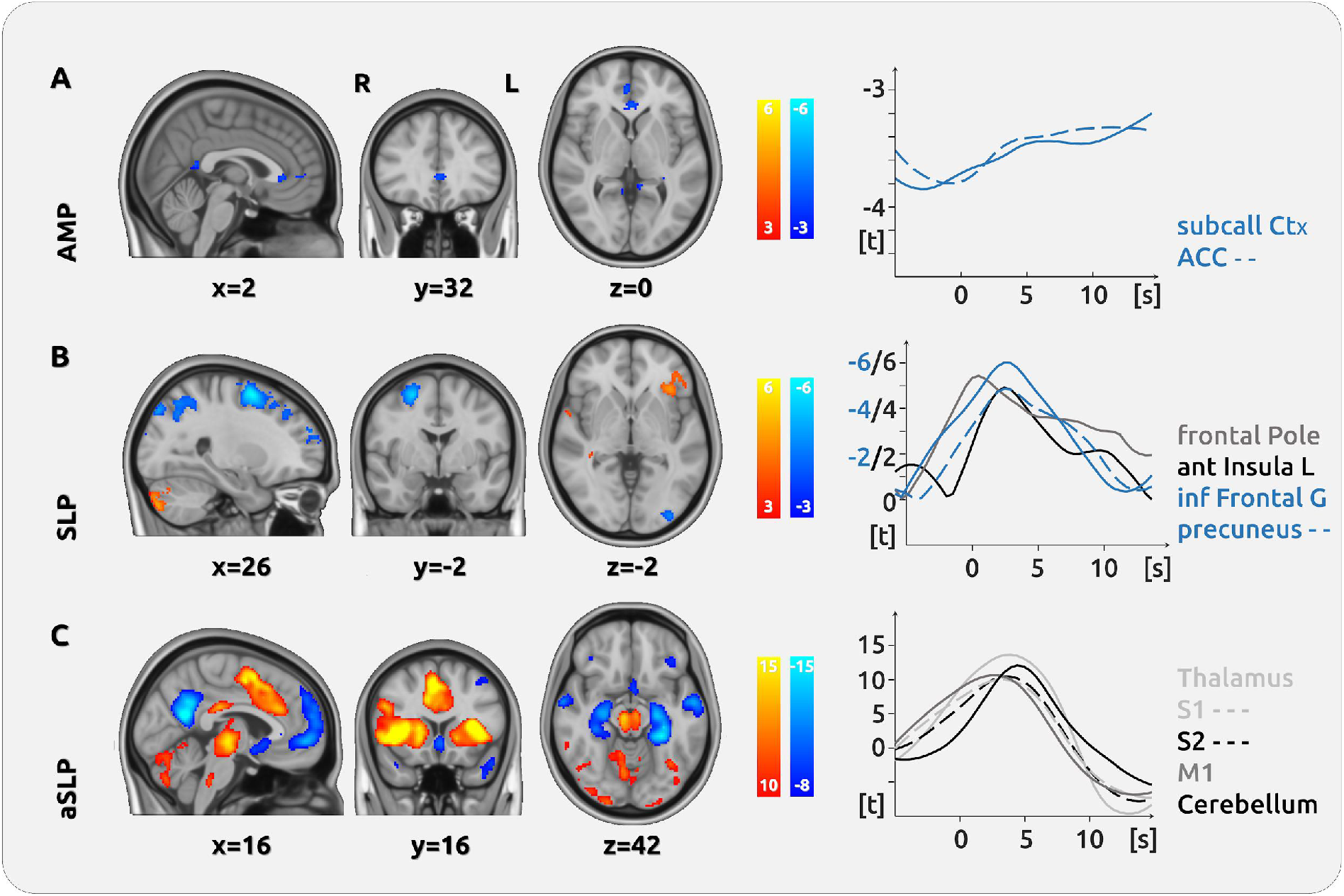
Cortical processing of chronic pain in CM. (A) The upper row shows no major region that encodes the intensity of endogenous pain (*amplitude* - AMP). We found negative relationships between brain activity and pain intensity in the PCC and the perigenual ACC. (B) The processing of changes of pain intensity (*slope* - SLP) was mainly localised in the left anterior insular cortex. Negative relationships were found in frontal and motor areas as well as in the precuneus. (C) The movement of the slider entails the activation of a large network of brain regions, which reflects motor activity and decision making (*absolute slope* - aSLP), shows a vast network of activity in the thalamus, the cingulate cortex, the entire insula, and the cerebellum. The graph (right) shows the temporal dynamics of the haemodynamic response for several regions in relation to a current pain rating at time point (= 0 s).

#### Encoding of the change of pain intensity across all CM patients (SLP)

The processes that contribute to the change of pain intensity activate the cerebellum, regions in the frontal lobe (frontal orbital cortex, inferior frontal gyrus), the lingual cortex, and the AIC. Brain regions that exhibit decreased activity during increasing pain comprise various regions in the occipital lobe (occipital pole, lateral occipital cortex, occipital fusiform cortex), as well as the middle frontal gyrus, the precuneus, and the pre- and postcentral gyrus (Fig. 4B; Table 6).

#### Processing of motor activity and decision making across all CM patients (aSLP)

The processing of motor activity, decision making, and changes of visual input shows a network of increased activity in the frontal operculum cortex, the insular cortex, the thalamus. Decreased activity was found in the precuneus, the PCC, the hippocampus, the lateral occipital cortex, the pACC, and the cerebellum. To separate the large overlapping clusters and to disentangle the contribution of the brain regions, we increased the cluster threshold beyond the threshold of the randomisation statistics. This affects the number of activated voxels; the figure has been generated based on the original PALM threshold (Fig. 4C; Supplementary Table 4).

#### Visual control experiment

In our control experiment, predominantly occipital areas (occipital pole, lateral occipital cortex and occipital fusiform gyrus) of the brain were activated for both patient groups, CBP and CM. We did not differentiate between the two patient groups for the analysis of the visual control experiment (see Supplementary Table 5).

#### Individual patterns of pain intensity encoding in single patients

Activation maps for intensity encoding (AMP) for all single patients are shown in Supplementary Material (Supplementary Fig. 3 for CBP and Supplementary Fig. 4 for CM). We found a considerable variety of activation patterns: while some patients exhibit a similar activity map as the maps from the group statistics, other patients show a rather weak correlation with the activity pattern of the group statistics (Fig. 5A). This variability is also reflected by the substantial individual pain-related activity of the AIC (Fig. 5B).

**Figure 5.**
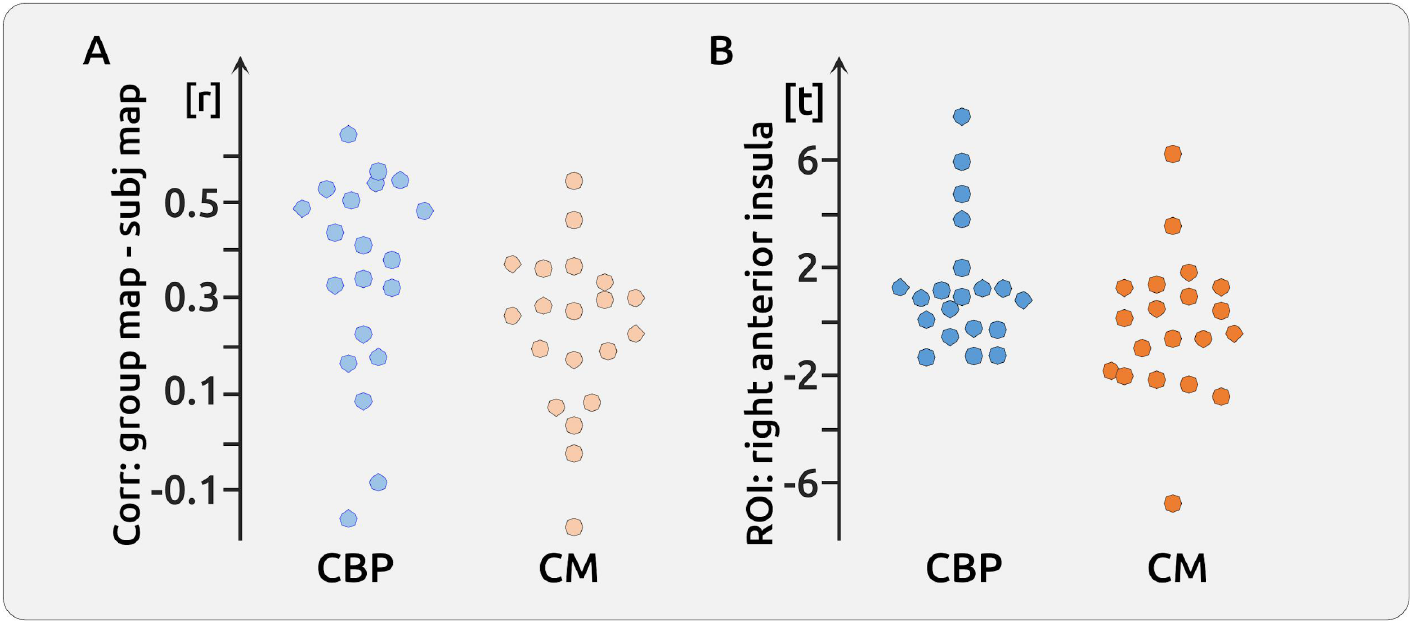
Individual patterns of pain intensity encoding. (A) Correlation of individual maps with the map based on group statistics revealed a correlation of 0.36 (±0.22) for CBP (left) and of 0.24 (±0.17) for CM (right) (B) Encoding of pain intensity in the right insula for CBP and CM: We found a correlation of 0.26 (±0.21) for CBP and of 0.21 (±0.24) for CM.

## Discussion

Here, we aimed to investigate how the intensity and intensity change of endogenous pain is subserved in the brain of chronic pain patients. The experimental design resembles the everyday experience of these patients, which is characterised by naturally-fluctuating pain and consists of phases of relatively low and periods of relatively high pain. Therefore, we are discussing the findings of the present study mainly in light of previous approaches to investigate endogenous and fluctuating chronic pain.

*First*, we have identified brain regions that are involved in the encoding of pain intensity irrespective of whether the pain is currently rising, falling or stable. *Second*, we detected brain regions that encode transient states of increasing and decreasing pain; these processes could be targeted in a future neurofeedback study with the aim to facilitate decreasing pain. *Third*, our statistical model included aspects that are not specific to pain processing, i.e. decision-making, visual change processing, and motor activity. This variable was included in the model in order to control for confounds in the relationship between the timecourse of AMP and cortical activity. It must be noted that motor activity, decision and visual processes can not be separated in the aSLP variable. *Fourth*, we emphasise the importance of a more patient-centred approach in neuroscience by repeatedly recording and analysing single patients. Consequently, each patient shows an unique pattern of cortical pain encoding, which can be considered as an individual signature of chronic pain processing. Many of the individual signatures only distantly resemble the result of the group statistics.

### Encoding of pain intensity across all patients - AMP

For CBP patients, we found - across all patients and sessions - that the intensity of pain is mainly encoded in the AIC, the frontal operculum, and the pons. These regions exhibit a positive relationship between cortical activity and the amplitude of continuous pain ratings: higher intensities of pain are accompanied by higher cortical activity. Additionally, we found that higher intensities of endogenous pain in CBP are related to decreased activity in the pACC, the precuneus, the hippocampus, and several occipital regions. This network of brain regions is largely overlapping with the NPS (Wager et al., 2013), but contradicts findings from previous work that located the encoding of *endogenous chronic* lower back pain predominantly in frontal regions (Baliki et al., 2006; Hashmi et al., 2012a, 2012b).

For the encoding of the pain intensity in CM, we did not observe any cortical region within the NPS network but found a positive relationship between pain intensity and cortical activity in the fusiform cortex. In addition, similar to the CBP patients, we found a negative relationship between pain intensity and brain activity in the pACC, the precuneus, and the hippocampus, as well as in the thalamus and the subcallosal cortex. The absence of any single region that codes for pain intensity in CM underlines the complex pathophysiology of the disease (Burstein et al., 2015; Dodick, 2018). In line with our initial hypothesis, we observed a complex and patient-specific pattern of the cortical processing of CM, which is not reflected in the group statistics; this is echoed in the individual profiles of endogenous pain processing in each CBP patient (see a more in-depth discussion on individual patterns below).

For the present investigation, we kept and analysed the full dynamics of the fluctuating endogenous pain perception of chronic pain patients; we did not define any statistical “events” such as periods of high pain or periods of increasing pain, but instead took the entire naturally-evolving trajectory of pain into account. We also applied a very long time constant for high-pass filtering the data to preserve the natural shape of the pain-rating timecourse. This preservation also excluded any direct correction for autocorrelation, which could have otherwise destroyed the natural structure of the data (see Methods section for a more suitable strategy regarding the correction of autocorrelation for continuous data). Furthermore, we did not include patients who would not exceed a certain amount of pain fluctuation, i.e. by excluding recordings with insufficient fluctuations or even steadily increasing pain. The extended duration of the experiment guaranteed balanced phases of increasing, stable, and decreasing pain, which disentangled the processes related to pain intensity from aspects of pain intensity changes in *all* patients (see single session pain rating trajectories in Supplementary Fig. 1 and Supplementary Fig. 2).

### Encoding of the change of pain intensity across all patients - SLP

Although the fluctuating pain intensity represents an important daily life experience of chronic pain patients, this phenomenon is barely investigated. Here, we explored how the processing of pain *intensity changes* is subserved in the brain and found that the change of endogenous pain of CBP patients is mainly encoded in the anterior insular cortex and in frontal regions. Regions that increase pain activity with falling pain were found in the lateral occipital cortex and the cerebellum. For CM patients, positive relations were located in the anterior insula, the hippocampus, the cerebellum, the orbitofrontal cortex, and the lingual gyrus. Besides occipital regions, negative relations were found in the middle frontal gyrus, the precuneus, the pACC, S1, M1 and the thalamus.

Similar to our results, Baliki and colleagues reported increased activity in the right anterior insula and the cerebellum, but showed additional activity in the posterior insula, S2, multiple portions of the middle cingulate cortex, and S1 (Baliki et al., 2006). However, these findings apply to periods of increasing pain contrasted to both periods of stable and decreasing pain. Therefore, the analysed periods of increasing pain are more influenced by pain-unspecific effects of motor activity and decision-making. In the present study, we have controlled for these aspects of motor activity and decision-making by contrasting time periods of rising pain with periods of falling pain. Due to further prerequisites of balanced and fluctuating pain ratings, the analysis of pain changes can be considered as independent from the analysis of pain amplitude: increasing pain can occur at both high and low levels of pain. The regions that encode the change of pain experience could have an impact on the emotional well-being of pain patients and would be a valuable target for therapeutic interventions such as neurofeedback.

### The role of the anterior insular for the processing of physical and chronic pain

For both pain diseases, we found the largest cortical effect for pain direction encoding (SLP) in the anterior insula; rising endogenous pain is accompanied by increasing insular activity. Interestingly, for CBP the AIC is processing the intensity (AMP) and the change of pain intensity (SLP). However, the encoding of intensity exhibits a different temporal profile and has its peak slightly before the processing change detection. Therefore, the activity of the AIC for amplitude and change of amplitude is overlapping and is determined by a summation of the activity for the current pain intensity and the transient activity for rising (plus) or falling (minus) pain. The results of the present investigation, in particular the contribution of the anterior insula, does not exclude that chronic pain patients suffer from emotional problems (Baliki et al., 2006; Hashmi et al., 2013), but militates against opinions that the transitions to chronic pain might be reflected by a shift from insular processes to frontal areas. Our findings argue against suggestions that chronic pain may have become decoupled from the sensory aspects but are purely emotional. These new findings could help to prevent the stigmatisation of pain patients as being mentally impaired (Cohen et al., 2011; De Ruddere and Craig, 2016).

### Individual patterns of pain intensity encoding in single patients

A major aspect of our investigation is the reliable assessment of a single patient’s profile of pain processing. In order to describe unique patterns of pain processing, each pain patient was recorded four times. As a result, the individual maps differ remarkably from each other and we found a substantial variation for the correlation of single patient signatures with the activity pattern from the groups statistics. For CBP, the activity of the prominent regions of the group statistics may have been driven by the few patients who exhibit strong effects in the anterior insula cortex and the ACC (see supplementary figures). As research articles usually do not publish single patient maps, we can only speculate whether this phenomenon can be generalised. Indeed, the importance of assessing individual patterns of brain structure and activity has been suggested previously (Martucci et al., 2014).

Studies that address single patient effects have reported some variability (Gordon et al., 2017; Greene et al., 2019; Marek et al., 2018). In a similar vein, and although group statistics suggest otherwise, individual parameters of gamma oscillations in tonic and chronic pain show that pain is not encoded by gamma activity in all study participants (May et al., 2019; Schulz et al., 2015). The enormous variability of the individual pain signatures indicates qualitative rather than quantitative differences between patients (Zadelaar et al., 2019). If this phenomenon would be true, this would suggest that the currently discussed “replication crisis” in neuroimaging would rather be a “sample crisis”, for which the replication of an effect would depend on whether the repeated study had included a similar “dominant” sub-sample of participants showing similar activity patterns. For these reasons we decided against a direct comparison of the brain processes of CM and CBP patient groups.

The different pain signatures in the present study depicts a more complex picture: even if all patients had a marginal (and positively correlated) contribution of the insular cortex (which they do not have), we must assume that the major weight for the encoding of pain for most patients relies on the processing in different brain regions. The variety of cortical processing is indeed in line with the clinical picture of pain diseases, in which each individual exhibits a complex composite of characteristics (Smith, 2009).

We can not exclude that some variables such as disease duration, psychological parameters, current medication, the current average pain intensity, as well as subgroups of the pain disease would modulate some aspects for a specific individual. However, the lack of significant and consistent positively correlated insular activity in most patients is suggested to be caused by qualitative rather than gradual differences between patients and would make the interpretation of quantitative effects challenging.

The question is, whether it is plausible that the encoding of pain in the insula (or any other region) can be modulated (e.g. by parameters like depression) in a way that some patients have a positive correlation between pain and brain activity and some patients would have negative relationships between brain activity and pain. The same applies for the pACC, which is considered to be a main hub of the descending pain control system. The group results suggest a negative relationship, but some individuals exhibit a positive relationship between the pACC and pain intensity, which would turn this region into a pain facilitation region rather than a pain inhibition hub. In other words, can some factors reverse the relationship between cortical activity and pain intensity? The plausibility of this potential phenomenon decides whether a correlation of any factor with the present variety of cortical maps is justified.

For the present study, we interpret the qualitatively different pattern as being modulated by other variables or subgroups as not plausible; to consider individual qualitative variation as noise might indeed limit our understanding on how fluctuations in pain over time are associated with pain processing, coping, and treatment response. Therefore, assessing individual pain signatures could facilitate more accurate assessment of chronic pain conditions, which is in line with recent developments to enhance and promote individually-tailored treatments in medicine (Mun et al., 2019; Ott et al., 2017).

## Limitations

The present investigation included a large sample of patients with repeated recordings. Although the entire data amounts to more than 100 minutes of pain encoding data for each single patient, there are a few limitations that need to be considered. The majority of patients were on different types of prescribed medication which could have had an effect on the BOLD fluctuations. In addition, even the four recordings for each patient might not be enough to assess the stable and invariant pain signature of an individual. Similarly, the statistics on single patients might have been driven by a subset of the recordings. The investigation on the stability of the individual pattern of pain-related cortical activity across the four recordings, however, would require a different methodological approach, i.e. machine learning tools, which is beyond the scope of the present study but will be addressed in a subsequent analysis.

## Conclusion

The present study expands the knowledge on the cortical underpinnings of chronic pain by showing that individuals exhibit their own signature of cortical processing of chronic pain; this applies to chronic back pain as well as to chronic migraine. This finding is matched by the experience of clinicians; each patient can be characterised by a unique personality with various combinations of symptoms and a vast range of treatment success, which is reflected by a wide range of individual responses to medical treatment. In the quest to find the optimal treatment for each pain patient, the findings support recent developments for a more personalised medicine. Consequently, the present findings argue against a common biomarker for the subjective experience of chronic pain that is based on neuroimaging.

On the contrary, as the experimental setup was aimed to reflect the unique and natural trajectory of the subjective experience of pain, the data would support an individually tailored therapeutic approach in clinical settings. Further studies are needed to explore whether the current findings on individual pain signatures can be utilised for interventions that aim to directly modulate brain activity, e.g. neurofeedback.

## Acknowledgements

We thank Dr Javeria Hashmi for her comments and Dr Stephanie Irving for copy-editing the manuscript. We thank Dr Tor Wager for his permission to use the NPS map.

