## Supplementary Material for "Chronic Pain Patients Exhibit Individually Unique Cortical Signatures of Pain"

**Supplementary Table 1 Characteristics of CBP patients reported in questionnaires**

| patient | m/f | age (years) | pain duration (years) | pain location | pain medication | pain intensity | PCS | d/a/s |
|---|---|---|---|---|---|---|---|---|
| 1 | f | 52 | 32 | thoracic | none | 4 | 14 | 1/4/11 |
| 2 | m | 64 | 2 | lumbar | none | 3 | 22 | 3/5/7 |
| 3 | f | 39 | 8 | lumbar | Ibuprofen 600 mg (10-12x/month)) | 4 | 32 | 7/9/16 |
| 4 | f | 41 | 18 | lumbar | Fluoxetine 20 mg (daily), Paracetamol 500 mg (2-3x/month), Orphenadrine 100mg (3x/month) | 3 | 33 | 6/7/11 |
| 5 | f | 60 | 14 | thoracic | Diclofenac 69,82 mg (1x/month) | 4 | 14 | 2/5/6 |
| 6 | f | 39 | 3 | lumbar | Paracetamol 500mg (2-3x/month), Ibuprofen 400mg (2x/month) | 4 | 13 | 2/1/1 |
| 7 | f | 48 | 5 | cervical | Ibuprofen 400mg (4x/month) | 5 | 12 | 4/0/6 |
| 8 | f | 52 | 11 | thoracic & lumbar | Ibuprofen 400mg (5x/month), Metamizole 500mg (2x/month) | 4 | 7 | 1/2/5 |
| 9 | f | 31 | 6 | lumbar | Ibuprofen 800mg (2-3x/month), Metamizole 1000mg (2-3x/month) | 5 | 1 | 0/0/1 |
| 10 | m | 26 | 4 | lumbar | none | 5 | 9 | 0/3/3 |
| 11 | f | 55 | 16 | thoracic & lumbar | Cannabis drops (15x/month) | 10 | 11 | 5/2/6 |
| 12 | m | 65 | 8 | lumbar | Ibuprofen 600mg (10-15x/month) | 4 | 10 | 2/2/3 |
| 13 | f | 31 | 1 | cervical & thoracic | Ibuprofen 400mg (18x/month) | 3 | 25 | 9/6/14 |
| 14 | m | 32 | 7 | thoracic and lumbar | Ibuprofen 600mg (6x/month), Tramadol 100mg (10x/month) | 5 | 22 | 7/3/5 |
| 15 | f | 26 | 10 | lumbar | Ibuprofen 400mg (1x/month) | 4 | 18 | 6/3/8 |
| 16 | f | 55 | 16 | cervical & thoracic | Metamizole 500mg (2-3x/month) | 5 | 2 | 4/0/4 |
| 17 | f | 56 | 15 | lumbar | none | 7 | 36 | 4/4/13 |
| 18 | f | 42 | 11 | thoracic & lumbar | none | 5 | 10 | 1/2/6 |
| 19 | f | 30 | 3 | thoracic | Ibuprofen 400mg (3x/month) | 4 | 27 | 2/4/8 |
| 20 | f | 43 | 10 | cervical & lumbar | Ibuprofen 400mg (5x/month) | 7 | 22 | 5/4/6 |

m/f: male/female; PCS: pain catastrophizing scale; d/a/s: depression/anxiety/stress. The cutoff for depression and stress is 10 and for anxiety 6.

**Supplementary Table 2 Characteristics of CM patients reported in questionnaires**

| patient | m/f | age (years) | pain duration (years) | pain medication | pain intensity | PCS | d/a/s |
|---|---|---|---|---|---|---|---|
| 1 | f | 61 | 50 | Sumatriptan 100mg (20x/month) | 7 | 15 | 0/10/9 |
| 2 | f | 27 | 7 | Metamizole 500mg (2-3x/month), Sumatriptan 50mg (1x/month) | 4 | 5 | 0/0/2 |
| 3 | f | 50 | 35 | Sumatriptan 100mg (5-7x/month) | 4 | 24 | 5/0/6 |
| 4 | f | 27 | 8 | Zolmitriptan 20mg (2x/month), Ibuprofen 600mg (7x/month) | 7 | 37 | 1/1/8 |
| 5 | m | 49 | 30 | Ibuprofen 600mg (7-8x/month), Metamizole 500mg (3-4x/month), Paracetamol 500mg (5-6x/month) | 4 | 3 | 0/6/1 |
| 6 | f | 52 | 30 | Ibuprofen 400mg (6x/month), Paracetamol 1000mg (2x/month) | 5 | 11 | 5/10/12 |
| 7 | f | 32 | 15 | Zolmitriptan 5mg (8x/month), Naproxen 500mg (15x/month), Acetylsalicylic Acid (ASA) 250mg (4x/month), Paracetamol 200mg (4x/month), Caffeine 50mg (4x/month) | 4 | 31 | 5/7/7 |
| 8 | f | 21 | 7 | Sumatriptan 50mg (1x/month) | 4 | 10 | 2/1/0 |
| 9 | f | 19 | 7 | none | 4 | 35 | 10/2/6 |
| 10 | f | 46 | 13 | Ibuprofen 800mg (8-10x/month) | 6 | 31 | 7/10/15 |
| 11 | f | 27 | 13 | Triptan (2-3x/month), ASA 250mg (20-25x/month), Paracetamol 250mg (20-25x/month), Caffeine 50mg (20-25x/month) | 4 | 13 | 6/1/10 |
| 12 | m | 53 | 15 | Ibuprofen 600mg (10x/month) | 6 | 20 | 7/2/4 |
| 13 | f | 30 | 6 | Ibuprofen 400mg (4x/month) | 4 | 24 | 1/1/5 |
| 14 | f | 21 | 7 | Ibuprofen 600mg (4-8x/month), Paracetamol 500mg (4x/month), Zolmitriptan 5mg (1-2x/month) | 3 | 15 | 2/0/5 |
| 15 | f | 23 | 8 | Ibuprofen 600mg (2-3x/month) | 7 | 24 | 2/0/3 |
| 16 | f | 28 | 7 | Ibuprofen 500mg (10x/month) | 7 | 11 | 0/1/1 |
| 17 | f | 25 | 5 | Ibuprofen 600mg (5x/month), Zolmitriptan 5mg (1x/month) | 4 | 21 | 5/5/9 |
| 18 | f | 33 | 20 | Paracetamol 500mg (3x/month) | 5 | 32 | 4/0/4 |
| 19 | f | 21 | 9 | Ibuprofen 400mg (6-10x/month), Rizatriptan 10mg (2x/month) | 5 | 30 | 1/1/2 |
| 20 | f | 43 | 10 | Ibuprofen 400mg (20x/month), Paracetamol 325 mg (8-10x/month), Naproxen 100 mg (8-10x/month), Caffeine 50 mg (8-10x/month), Drotaverine hydrochloride 40 mg (8-10x/month), Pheniramine 10 mg (8-10x/month) | 4 | 22 | 2/3/9 |

m/f: male/female; PCS: pain catastrophizing scale; d/a/s: depression/anxiety/stress. The cutoff for depression and stress is 10 and for anxiety 6.

**Supplementary Figure 1 Time courses of pain ratings for CBP patients**

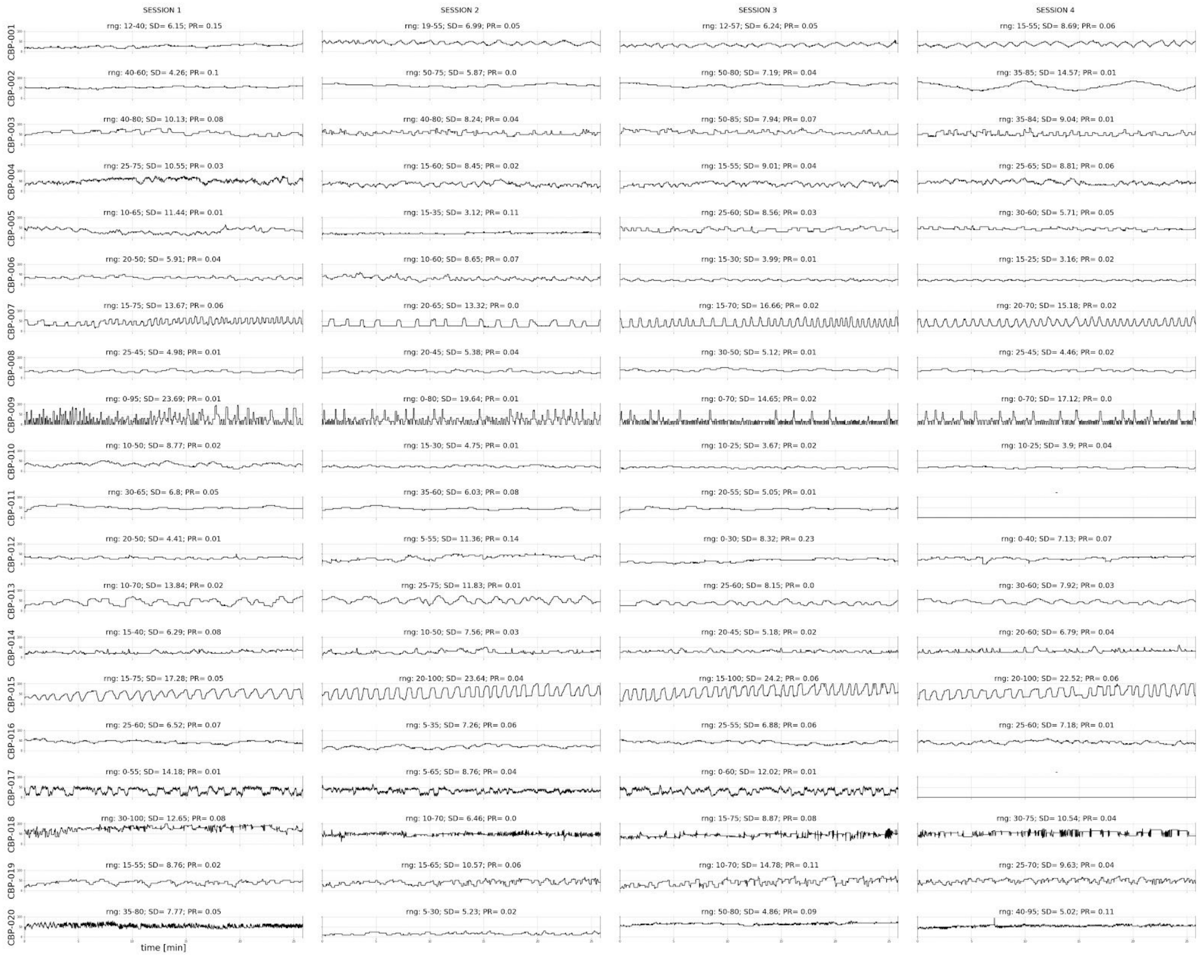


For all patients (except CBP-011 and CBP-017), four sessions of approximately 25 min were recorded successfully. The range (rng) of the pain intensity rating and the standard deviation (SD) over the whole time series are given. The ratings of each subject's pain was measured with the parameter PR to ensure a large variability in pain ratings, as well as a small to moderate increase of the overall pain ratings across the rating task.

**Supplementary Figure 2 Time courses of pain ratings for CM patients**

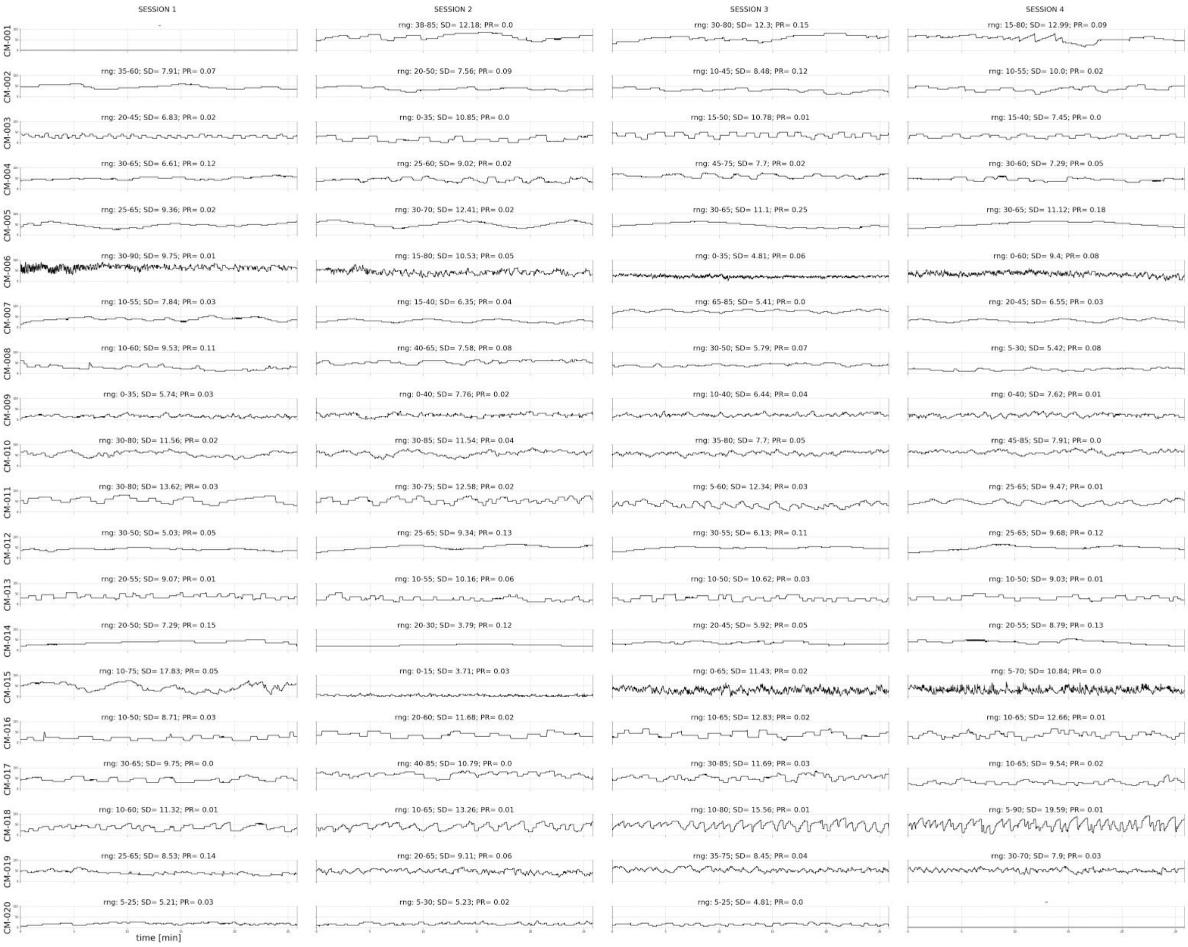


For all patients (except CM-001 and CM-020), 4 sessions of approximately 25 min were recorded successfully. The range (rng) of the pain intensity rating and the standard deviation (SD) over the whole time series are given. The ratings of each subject’s pain was measured with the parameter PR to ensure a large variability in pain ratings, as well as a small to moderate increase of the overall pain ratings across the rating task.

**Supplementary Figure 3 CBP, endogenous pain encoding of single subjects**

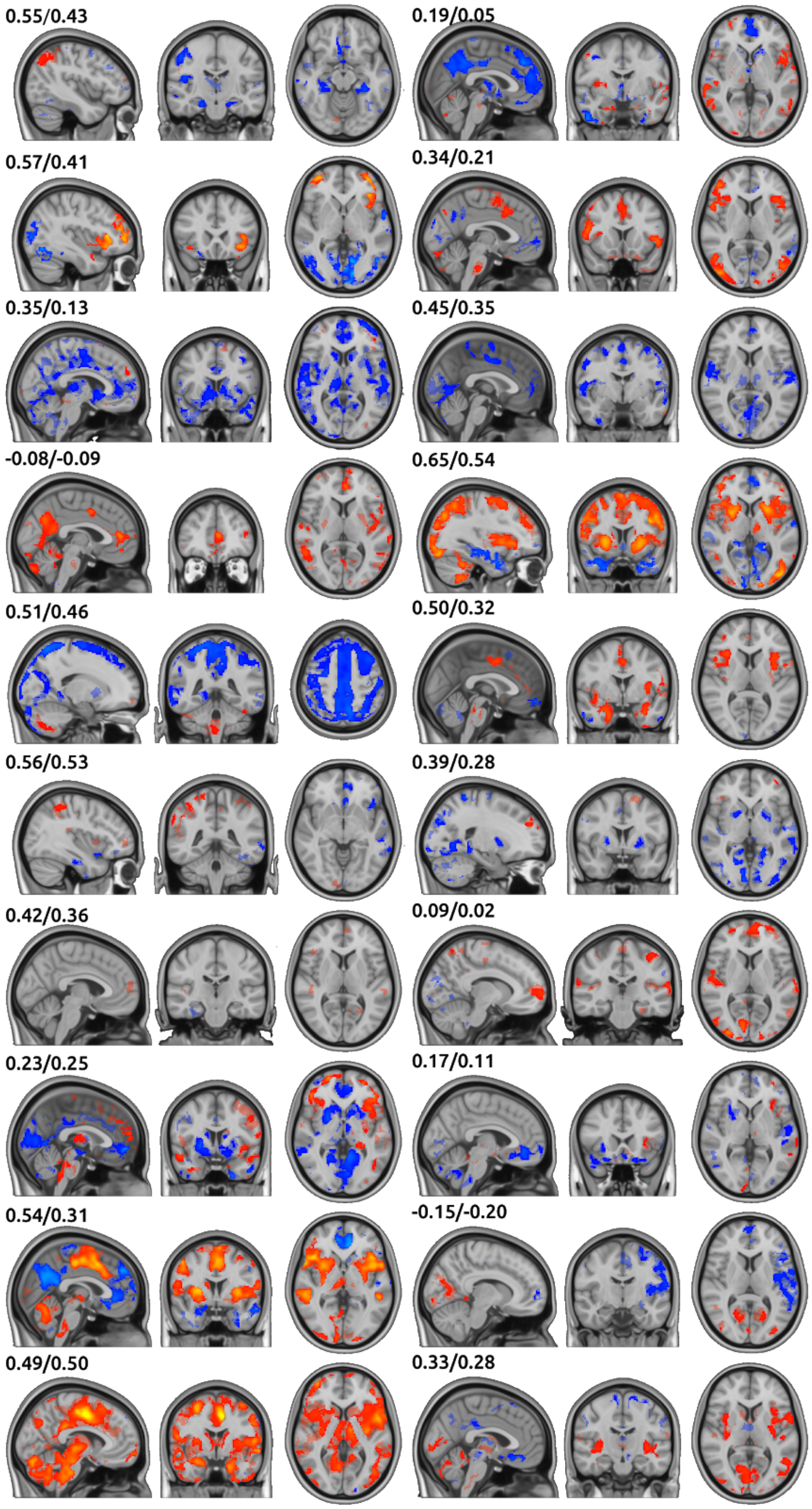

The numbers indicate the spatial correlation of the individual map with the group map (left number: correlation within the activity maps of the group statistics (t>2) / right number: correlation within the areas activated by the subject (t>2).

### Supplementary Figure 4 CM, endogenous pain encoding of single subjects

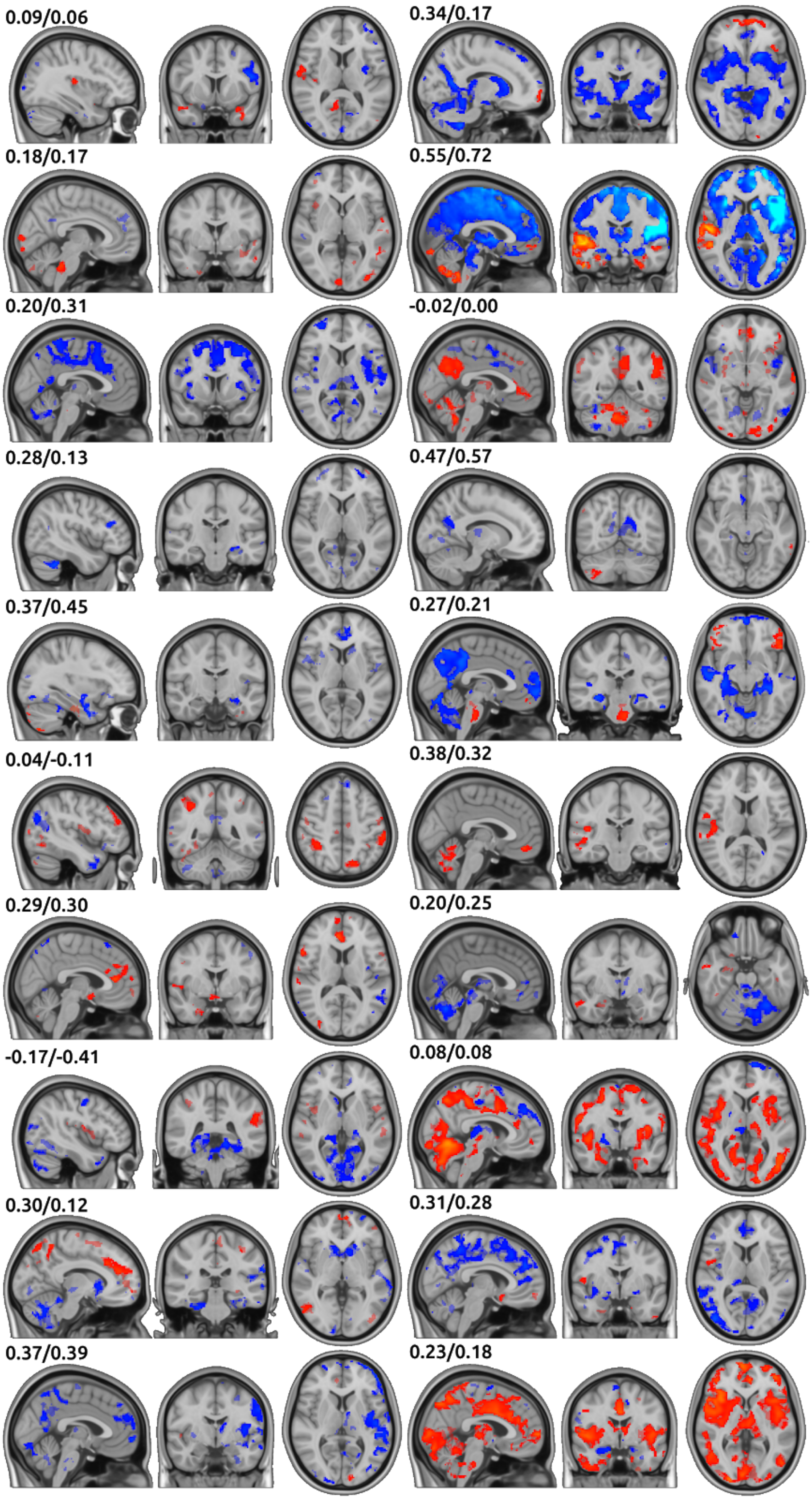

The numbers indicate the spatial correlation of the individual map with the group map (left number: correlation within the activity maps of the group statistics (t>2) / right number: correlation within the areas activated by the subject (t>2).

**Supplementary Table 3 Active brain areas that encode processing of motor activity, decision making, and visual change across all CBP patients**

| Anatomical structure | Cluster size | t-value | | Coordinates | | |
|---|---|---|---|---|---|---|
| | | pos | neg | x | y | z |
| Perigenual Anterior Cingulate Cortex (pACC) | 1108 | 16.1 | | -0 | 12 | 44 |
| Superior Parietal Lobe, Supramarginal Gyrus | 1065 | 19.5 | | -40 | -42 | 47 |
| Supramarginal Gyrus, Angular Gyrus | 1007 | 16.1 | | 44 | -43 | 46 |
| Inferior Frontal Gyrus, Precentral Gyrus (M1) | 921 | 16.7 | | 47 | 8 | 18 |
| Occipital Fusiform Gyrus, Lateral Occipital Cortex | 890 | 15.9 | | -31 | -79 | -15 |
| Hippocampus | 601 | | -12.8 | -27 | -31 | -13 |
| Insular Cortex | 582 | 15 | | -37 | 8 | 5 |
| Cerebellum (Right VI) | 566 | 16.6 | | 32 | -56 | -26 |
| Superior Temporal Gyrus, Middle Temporal Gyrus | 507 | | -11.8 | -56 | -10 | -9 |
| Hippocampus | 507 | | -10.9 | 29 | -26 | -14 |
| Lateral Occipital Cortex | 474 | | -11.1 | -45 | -70 | 34 |
| Precentral Gyrus (M1), Middle Frontal Gyrus | 444 | 17.1 | | -32 | -7 | 56 |
| Precuneus | 411 | | -11.1 | -8 | -59 | 21 |
| Occipital Pole | 379 | 17.3 | | 29 | -94 | 0 |
| Superior Temporal Gyrus, Middle Temporal Gyrus | 307 | | -10.2 | 60 | -11 | -7 |
| Thalamus | 257 | 15.5 | | -12 | -17 | 8 |
| Lateral Occipital Cortex | 213 | | -11.5 | 52 | -65 | 31 |
| Thalamus | 209 | 14.8 | | 12 | -15 | 8 |
| Perigenual Anterior Cingulate Cortex (pACC) | 195 | | -9.27 | -3 | 49 | -2 |
| Precuneus | 172 | | -9.28 | 8 | -58 | 22 |
| Central Opercular Cortex, Postcentral Gyrus | 170 | 16.7 | | -54 | -21 | 20 |
| Cerebellum (Right VIIIa) | 144 | 15.6 | | 24 | -58 | -48 |
| Frontal Pole, Middle Frontal Gyrus | 126 | 15 | | 38 | 36 | 32 |
| Occipital Fusiform Gyrus | 88 | 15 | | -39 | -65 | -13 |
| Occipital Fusiform Gyrus | 85 | 15 | | 40 | -63 | -13 |
| Subcallosal Cortex | 83 | | -9.41 | -0 | 10 | -8 |
| Temporal Pole | 57 | | -8.69 | -45 | 14 | -28 |
| Precentral Gyrus (M1) | 55 | 13.9 | | -50 | 3 | 32 |
| Posterior Cingulate Cortex (PCC) | 48 | 13.4 | | 1 | -29 | 26 |
| Frontal Pole, Frontal Orbital Cortex | 32 | | -9.04 | 38 | 35 | -10 |
| Frontal Orbital Cortex , Frontal Pole | 24 | | -9.21 | -39 | 33 | -11 |

| | | | | | | |
|---|---|---|---|---|---|---|
| Frontal Pole, Superior Frontal Gyrus | 13 | | -8.35 | -3 | 59 | 29 |

**Supplementary Table 4 Active brain areas that encode processing of motor activity, decision making, and visual change across all CM patients**

| Anatomical structure | Cluster size | t-value | | Coordinates | | |
|---|---|---|---|---|---|---|
| | | pos | neg | x | y | z |
| Perigenual Anterior Cingulate Cortex (pACC) | 2148 | | -13 | -7 | 50 | 21 |
| Precuneus, Posterior Cingulate Cortex (PCC) | 1789 | | -17.3 | -4 | -56 | 25 |
| Lateral Occipital Cortex | 1147 | | -15.2 | -43 | -70 | 35 |
| Hippocampus | 1089 | | -15.5 | -26 | -26 | -15 |
| Middle Temporal Gyrus, Superior Temporal Gyrus | 821 | | -12.5 | -54 | -1 | -18 |
| Hippocampus | 762 | | -13.3 | 27 | -19 | -15 |
| Frontal Operculum Cortex, Central Opercular Cortex, Insular Cortex | 739 | 17.9 | | 41 | 13 | 6 |
| Perigenual Anterior Cingulate Cortex (pACC) | 538 | 17 | | 5 | 14 | 45 |
| Inferior Frontal Gyrus | 405 | | -11.9 | -47 | 31 | -0 |
| Thalamus | 394 | 17.1 | | 12 | -11 | 4 |
| Middle Temporal Gyrus, Superior Temporal Gyrus | 344 | | -12.3 | 59 | -1 | -16 |
| Lateral Occipital Cortex | 336 | | -12.1 | 49 | -63 | 31 |
| Frontal Operculum Cortex, Insular Cortex | 318 | 17.9 | | -35 | 15 | 6 |
| Subcallosal Cortex | 263 | | -11.6 | -0 | 11 | -8 |
| Supramarginal Gyrus, Angular Gyrus | 250 | 16.2 | | 44 | -45 | 45 |
| Middle Temporal Gyrus, Superior Temporal Gyrus | 219 | | -10.5 | -55 | -36 | 0 |
| Cerebellum (Right Crus II) | 153 | | -11 | 28 | -81 | -36 |
| Frontal Pole | 116 | 16.3 | | 37 | 42 | 26 |
| Putamen | 85 | 15.7 | | 19 | 11 | 4 |
| Superior Temporal Gyrus | 78 | | -9.69 | 65 | -23 | 6 |
| Pallidum | 71 | 15.3 | | -15 | -3 | 2 |
| Middle Frontal Gyrus | 68 | | -9.17 | -40 | 16 | 52 |
| Thalamus | 56 | 15.2 | | -6 | -18 | -3 |
| Precentral Gyrus (M1) | 39 | 15.1 | | 37 | -5 | 57 |
| Cerebellum (Right IX) | 39 | | -10.1 | 5 | -51 | -46 |
| Frontal Pole, Frontal Orbital Cortex | 29 | | -9.84 | 37 | 36 | -11 |
| Superior Parietal Lobe, Supramarginal Gyrus | 26 | 15 | | -32 | -48 | 45 |
| Precentral Gyrus (M1), Middle Frontal Gyrus | 23 | 15.3 | | 45 | 3 | 33 |
| Cerebellum (Right VI) | 23 | 14.5 | | 29 | -56 | -21 |
| Temporal Pole | 20 | | -8.55 | 40 | 13 | -25 |
| Hippocampus | 3 | | -8.19 | 21 | -41 | 5 |
| Precentral Gyrus (M1), Postcentral Gyrus (S1) | 1 | | -8.1 | 50 | -10 | 36 |

**Supplementary Table 5 Active brain areas during the visual control experiment for AMP, SLP and aSLP**

| Anatomical structure | Cluster size | t-value | | Coordinates | | |
|---|---|---|---|---|---|---|
| | | pos | neg | x | y | z |
| **AMP** | | | | | | |
| Occipital Pole | 98 | | -5.83 | 16 | -90 | 31 |
| **SLP** | | | | | | |
| Occipital Pole | 696 | 6.89 | | 25 | -92 | -4 |
| Occipital Pole | 325 | | -5.48 | 14 | -92 | 23 |
| Occipital Fusiform Gyrus, Lingual Gyrus | 65 | | -4.78 | -18 | -76 | -9 |
| Postcentral Gyrus (S1), Precentral Gyrus (M1) | 46 | | -4.46 | -22 | -32 | 74 |
| Lateral Occipital Cortex | 26 | 4.04 | | -15 | -75 | 53 |
| Occipital Pole | 26 | | -3.94 | -20 | -100 | 10 |
| Lateral Occipital Cortex | 23 | | -4.2 | -41 | -73 | 8 |
| Lateral Occipital Cortex | 17 | 3.31 | | -49 | -73 | 7 |
| Occipital Pole | 6 | 0.643 | | 15 | -96 | 26 |
| Occipital Pole | 5 | 2.8 | | -16 | -101 | 18 |
| Lateral Occipital Cortex | 4 | | -3.46 | -47 | -79 | 10 |
| Lateral Occipital Cortex | 1 | 1.36 | | -44 | -72 | 2 |
| Occipital Pole | 1 | 1.17 | | 16 | -94 | 12 |
| Occipital Pole | 1 | 0.688 | | 16 | -98 | 14 |
| **aSLP** | | | | | | |
| Posterior Cingulate Gyrus, Precuneus | 22270 | 17.6 | | 4 | -50 | 9 |
| ACC, pACC | 2124 | | -13.6 | -2 | 43 | 2 |
| PCC, precuneus | 1050 | | -14.9 | -3 | -53 | 28 |
| Lateral Occipital Cortex | 769 | | -14 | -45 | -66 | 37 |
| Thalamus | 642 | 14 | | -14 | -12 | 9 |
| Hippocampus | 378 | | -11.6 | -25 | -21 | -14 |
| Frontal Operculum Cortex, Central Opercular Cortex, Insular Cortex | 356 | 12.7 | | -40 | 12 | 5 |
| Lateral Occipital Cortex | 236 | | -11.9 | 53 | -62 | 33 |
| Frontal Orbital Cortex, Frontal Pole | 95 | | -11.2 | -35 | 35 | -10 |
| Hippocampus | 92 | | -10.8 | 25 | -15 | -16 |
| Middle Temporal Gyrus, Superior Temporal Gyrus | 83 | | -9.77 | -59 | -11 | -8 |
| Cerebellum (Left IX, Left VIIIb) | 50 | 11.1 | | -12 | -55 | -49 |
| Temporal Pole | 45 | 8.95 | | -43 | 14 | -31 |
| Superior Temporal Gyrus | 32 | | -9.73 | 64 | -26 | 8 |
| Central Opercular Cortex | 31 | | -9.08 | 58 | -8 | 8 |
| Superior Frontal Gyrus | 28 | | -8.88 | -21 | 32 | 51 |

| | | | | | | |
|---|---|---|---|---|---|---|
| Inferior Frontal Gyrus, Middle Frontal Gyrus | 25 | 10.2 | | -49 | 26 | 21 |
| Frontal Pole | 5 | | -8.55 | -12 | 42 | 49 |
| Central Opercular Cortex, Insular Cortex | 5 | | -8.15 | 38 | -15 | 19 |
| Supramarginal Gyrus | 2 | 9.36 | | 64 | -40 | 37 |
| Supramarginal Gyrus, Angular Gyrus | 1 | 8.57 | | 64 | -46 | 32 |
| Middle Frontal Gyrus | 1 | 8.53 | | 50 | 20 | 40 |
